# Increasing evenness and stability in synthetic microbial consortia

**DOI:** 10.1101/2022.05.25.493411

**Authors:** Ruhi Choudhary, Radhakrishnan Mahadevan

## Abstract

Construction of successful synthetic microbial consortia will harbour a new era in the field of agriculture, bioremediation, and human health. Engineering communities is a complex, multi-dimensional problem with several considerations ranging from the choice of consortia members and spatial factors to genetic circuit performances. There has been a growing number of computational strategies to aid in synthetic microbial consortia design, but a framework to optimize communities for two essential properties, evenness and stability, is missing. We investigated how the structure of different social interactions (cooperation, competition, and predation) in quorum-sensing based circuits impacts robustness of synthetic microbial communities and specifically affected evenness and stability. Our proposed work predicts engineering targets and computes their operating ranges to maximize the probability of synthetic microbial consortia to have high evenness and high stability. Our exhaustive pipeline for rapid and thorough analysis of large and complex parametric spaces further allowed us to dissect the relationship between evenness and stability for different social interactions. Our results showed that in cooperation, the speed at which species stabilizes is unrelated to evenness, however the region of stability increases with evenness. The opposite effect was noted for competition, where evenness and stable regions are negatively correlated. In both competition and predation, the system takes significantly longer to stabilize following a perturbation in uneven microbial conditions. We believe our study takes us one step closer to resolving the pivotal debate of evenness-stability relationship in ecology and has contributed to computational design of synthetic microbial communities by optimizing for previously unaddressed properties allowing for more accurate and streamlined ecological engineering.

## Introduction

Traditionally, single microbes have been engineered to incorporate an array of desired behaviours. However in recent years, microbial consortia have drawn significant attention. Engineering microbial consortia offer several advantages over engineering monocultures. By dividing the genetic processes between different populations the metabolic burden can be distributed, counteracting growth defects observed in engineered monocultures where the entire load is imposed on the single species ^1^. The diverse genetic arsenal of multiple populations can be harnessed to expand the spectrum of novel applications, allowing implementation of complex pathways. A major drawback with engineered monocultures is that non-orthogonal pathways can lead to unforeseen interactions. This undesired effect can be mitigated by allocating reactions to different populations ^2^. Additionally, since microbial communities more commonly occur in nature as opposed to isolated microbes, making an accurate investigation of consortia and ecological dynamics essential. Therefore, synthetic microbial consortia have emerged as the next frontier genetic manipulation for biotechnological purposes ^3, 4^. Their applications are widespread and not limited to chemical production^5, 6^, bioremediation ^7^, agriculture ^8^, and even tackling climate change ^9^.

Designing synthetic microbial consortia requires a mechanistic understanding of cellular social interactions and their relationship with two most essential ecological factors: diversity and stability. Diversity has been positively linked with bio-productivity in communities ^10^. Furthermore, diverse systems have a survival advantage, as in case of an invasion or obstruction leading to death of one species, the other species in the system can take over. Biodiversity of a community has two components: richness and evenness. The former represents the number of species present in a community while the latter measures the proportional abundance of species. Low values of microbial evenness indicates that one or few species dominate the community and high values indicate an equitable distribution of species. Even though microbial evenness has been known to influence community dynamics^11, 12, 13^, it remains a relatively understudied concept compared to species richness. High evenness has been shown to be desirable for enhancing community productivity by amplifying the functionality of each species ^14^ and for being more resilient to disturbances and stresses ^15^. Stability is an important metric indicative of the long-term performance and survival of a community. A microbial community can exhibit multiple stable states in response to invaders and other environmental factors ^16, 17^. Therefore when optimizing the performance of synthetic microbial consortia, it is ideal to ensure both high evenness and high stability. This is a multi-layered challenge. Firstly, the relationship between evenness and stability has been conflicting and not straightforward ^18^, and secondly, the system design by intuition becomes increasingly difficult as the size and complexity of the consortia grows. While stability has been studied in microbial communities ^19^, to our knowledge no one has engineered stability in microbial communities. Even though population evenness was shown to increase by engineering cross-feeding in a synthetic microbial community^20^, this increase in evenness was not coupled with stability.

To aid in fast and accurate construction of synthetic microbial communities, research groups have developed a range of computational methodologies^21, 22, 23, 24^. A network-based model CoMiDa was developed to identify minimal number of species using integer linear programming that could provide enough enzyme capacity for the production of desired metabolites from pre-defined range of substrates ^25^. MultiPus is another approach that employs the same principles to minimize the number of reactions in a community^26^. Several flux balance analysis (FBA) based strategies have emerged that mathematically model community behaviour and community *in silico* design ^27, 28, 29^. A multiscale framework, CODY (COmputing the DYnamics of the gut microbiota), uses spatio-temporal modeling to accurately quantify variations in species abundances in human colon ^30^. Another notable example is FLYCOP (FLexible sYnthetic Consortium Optimization) which predicts the optimized community configuration by taking into account the growth medium composition and temporal changes in metabolite concentrations^31^. Klamt *et al* developed a computational tool, ASTHERISC (Algorithmic Search of THERmodynamic advantages in Single-species Communities), that designs communities to maximize the thermodynamic driving force for product synthesis^32^. A group explored two- and three-strain systems using Bayesian methods and identified robust community configurations for steady state behaviour ^33^. Venturelli *et al* designed an approach to scan community design space for high-butyrate producing community assemblies^34^. A toolbox called MDSINE was developed by Bucci *et al* to infer interactions between species in a community and predict future dynamics^35^. Another group used a genetic algorithm to select optimal environmental composition to produce target community phenotype ^22^. While the established computational methodologies have made tremendous contribution to the construction of synthetic microbial consortia, none of these address the design process optimizing for microbial evenness and stability, two key factors that determine the success of a microbial community.

The widespread approach has been constructing optimal configuration by selecting social interactions and network motifs. Contrary to other studies, our work focuses on developing and applying a computational strategy to existing configurations to modulate evenness and stability. Intelligent manipulation of synthetic ecological models requires a holistic understanding of the system and the circuit parameters involved. Our study proposes a computational framework to identify the circuit parameters and the corresponding values to balance evenness and stability. Our workflow combines machine learning and mathematical modeling to explore and characterize the parametric space of circuit parameters to recalibrate their values to achieve an even and stable model. With this framework, we aim to gain crucial insights into social interactions and the circuit parameters contributing to evenness and stability. Using this method, we derive more streamlined experimental decisions to optimize synthetic microbial in a swift and accurate manner.

## Results

### Computational framework to explore and design synthetic microbial consortia

Synthetic microbial ecosystems consist of multiple species and several interactions in order to accomplish innovative tasks. However the first step in our understanding of them is to break them down into smaller and more digestible units. For that purpose, we created three interaction models of co-cultures - the simplest unit of a microbial consortia. Our goal was to examine social interactions in context of evenness and stability at the most fundamental level. However, our methodology can easily be modified to incorporate more species and relationships.

In our study, we have chosen competition, cooperation, and predation as test cases, as they are among the most fundamental interactions found in nature and heavily influence community dynamics^36^. Interactions such as cooperation can be beneficial to biotechnology purposes such as producing a product of interest^37^. Competition is more commonly found in nature, hence essential for understanding natural systems^38^. Predator-prey interaction is evolutionary important^39^ and has been shown to result in oscillatory or bistable behaviour ^40^. To simulate them, we targeted the production of two amino acids (tryptophan and histidine) using quorum-sensing (QS) system. QS-based circuits are essential for cell to cell communication and are obvious targets for ecological engineering^41, 42^. Figure 1A shows the three circuits of *E*.*coli* co-cultures. Species 1 and Species 2 are auxotrophic for tryptophan and histidine respectively. Small quantities of amino acids are added in the extra-cellular compartment to commence growth, after the consumption of which the species must rely on their cellular dynamics. As tryptophan user grows, AHL1 molecules accumulate. When a high enough cell density is reached, the AHL1 molecules propagate to the other *E*.*coli* and bind to LuxR protein causing a structural change and hence activating it. The complex induces target histidine production gene regulation in histidine user. Similarly, the growth of histidine user, leads to AHL2 molecules accumulation, ultimately regulating tryptophan production gene in the other species. The parameters involved in our systems are listed in Figure 1B. These have been identified as parameters that can be engineered in a wet-lab setting and play a role in cellular dynamics. We are interested in finding the optimal operating ranges for the parameters as opposed to precise values, which can be calculated using an optimizer, due to two reasons: first, biological systems are inherently noisy with variability arising from both intrinsic and extrinsic factors ^43^ and second, to account for ease of engineering. The ranges we chose for each parameter to explore within can be found in Methods.

**Figure 1:**
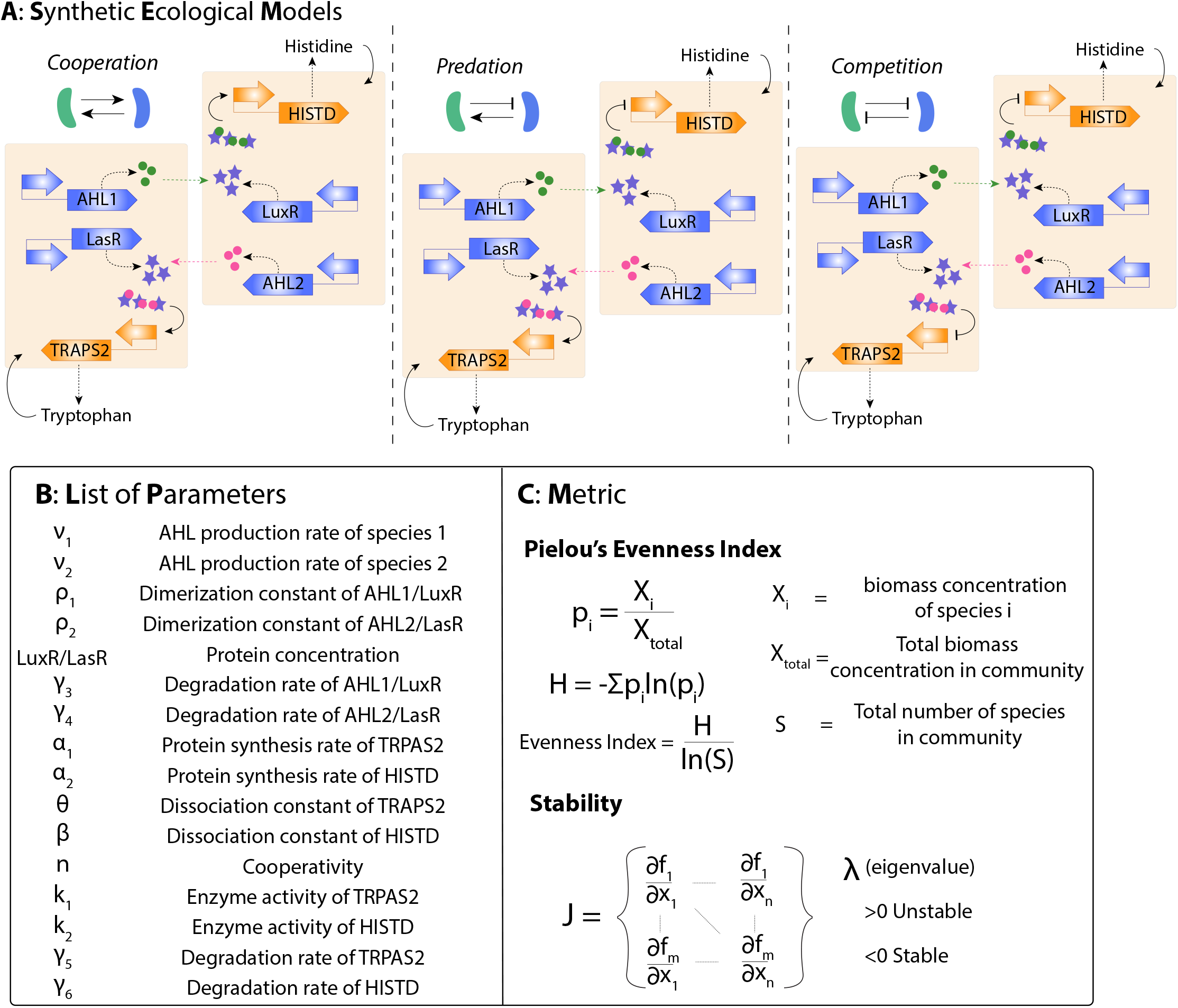
A) Quorum-sensing based circuits of cooperation, predation, and competition. Two E.coli species are auxotrophic for tryptophan and histidine respectively and depend on each other for survival. In short, the growth of one species triggers or suppresses the essential amino acid production in the other species depending on the type of circuit interaction. B) A list of tunable circuit variables that are involved in the synthetic models described above. The design space of these parameters were explored and characterized by our methodology. C) The two outputs evaluated are evenness and stability. The evenness index used is the Pielou’s evenness measure. Stability was calculated using linear stability analysis and eigenvalues with real negative eigenvalues denoting stable system.

Genome-scale modeling has provided valuable insights in the field of metabolic engineering and has found application in microbial community studies as well. To simulate our synthetic consortia models, we used a python version of dynamic FBA-based methodology called DMMM (dynamic multi-species metabolic modeling), which predicts growth dynamics of multiple species utilizing a single substrate source, as the foundation to develop our proposed framework ^44^. We modeled two species of *E*.*coli* - tryptophan user and histidine user-using a system of ordinary differential equations found in Methods section. We assumed a perfect chemostat to be able to achieve a steady state. A steady flow rate containing glucose was supplied to the system and the same flow rate exited the system, accounting for cell death. The ranges tested for circuit parameters are also listed in Methods. This led to a wide array of parametric combinations, significant portion of which resulted in washout. To mitigate this, we fixed the flow rate to a very low level to ensure that majority of our simulations reached a steady state. We defined steady state when change in biomass for at least three hours was minimal.

Given that seventeen parameters results in a very high dimensional space, the exploration of this parametric space can be very challenging. To cover the entire space satisfactorily, the number of simulations required increases in a combinatorial manner with the number of parameters and total operating ranges of the parameters^45^. For p number of parameters and n number of points sampled for each parameter, the minimum number of simulations needed would be n^p^. In our case, we have 17 parameters, even if we were to sample just 10 points per parameter, we would need to conduct at least 10^17^ simulations. This computationally expensive process poses a significant barrier in our ability to investigate the parametric space and draw useful information from it. We have developed methodology that enables swifter exploration and analysis of parametric space conceptualized in Figure 2. Since the biggest obstacle in mapping parametric space is its complexity and vastness, and hence the number of simulations required to accomplish it, the core principle of our methodology is to first reduce the parametric space without losing any critical information.

**Figure 2:**
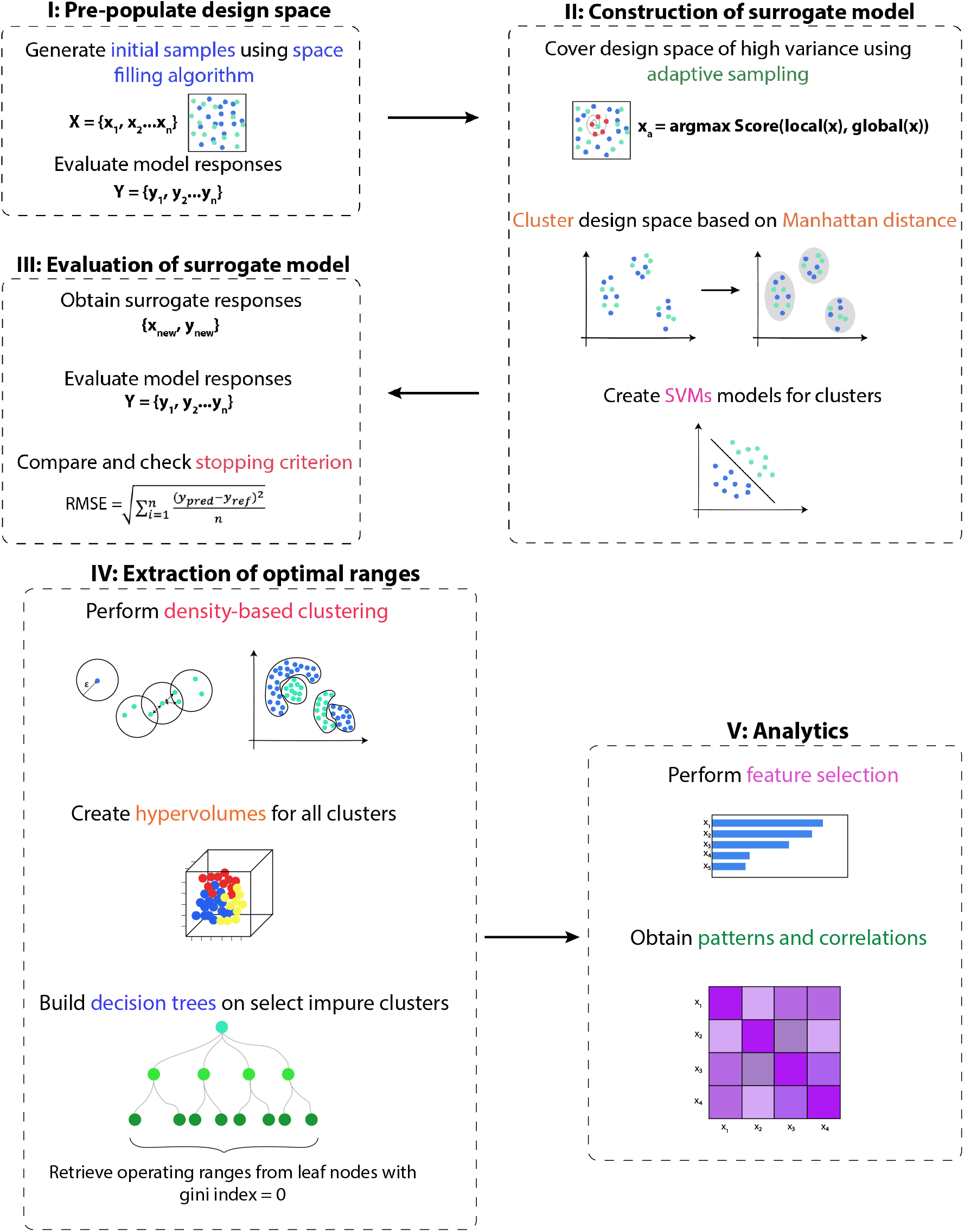
Machine learning based workflow can be divided into two parts. In the first section we use adaptive sampling to build surrogate models of our test cases to ensure that parametric space has been covered efficiently. The first step is to initially populate the space with Sobol points, following which adaptive sampling covers the design space by sampling in area of high variance and using previous points to select new points. To evaluate the efficient coverage of design space, surrogate models were constructed and their accuracy was determined using support vector machine algorithm. In the second section, we analyzed the parametric space using the principles of clustering and hypervolume to derive specific parameter ranges corresponding to our desired output. Once there are enough points in the parametric space, density-based clustering is applied to partition the space. Hypervolumes are created around each clusters, which provide their corresponding parametric ranges. To obtain optimal ranges, decision trees are used. Finally, further analysis such as feature importance can be performed.

Complex simulation models are often treated as “black-box” functions with unclear input-output mapping. They also incur high computational costs, which has led to the emergence of surrogate models, also known as metamodels ^46^. They capture the essential properties of the original model, emulating the model’s behaviour while mitigating computational costs. They are often used to replace the computationally expensive task with approximate functions that are faster and easier to evaluate. The first step in surrogate model construction is evaluating the original model at training points. To kickoff this process, we generated Sobol sequence numbers to generate an initial set of training points. Sobol numbers are space-filling and widely used. Once an initial set of points were obtained, we switched to adaptive sampling method to select next training points in the solution space. Adaptive sampling technique generates points in regions of the parametric space with large prediction errors ^47, 48^. Surrogates built from adaptive training points are better approximations while requiring fewer points and hence evaluations. It is a sequential sampling technique that draws upon the previous point to sample the next point and covers more space with fewer points compared to Sobol. To evaluate if the parametric space has been effectively covered, we tested the accuracy of the surrogate model. However since the parametric space is large and high dimensional, first we clustered the points generated based on Manhattan distance. Then we applied a supervised machine learning algorithm, support vector machine (SVM), on each cluster. Therefore we essentially tested the coverage by testing multiple surrogate model responses. Once the error was stabilized and below a threshold, we concluded that the design space has been covered and analysis can begin.

The design space has been populated with points from both sobol and adaptive sampling. For each point we calculated the evenness, based on the equation in Figure 1C.

As several clusters are generated, we had to select the cluster that is more suitable for experimental implementation. A combination of two features guide cluster selection: “probability” and “hypervolume”. The probability of a cluster describes the number of points inside it that correspond to desired output, with a higher probability being preferable. The hypervolume measures the operating ranges of the parameters. Depending on experimental constraints, specific values of parameters might be more suitable. To handle impure clusters, which mean clusters containing a mixed bag of high evenness and low evenness points, we built decision trees on them. Decision trees identified the parameter(s) within the cluster that required further tuning. In decision trees, Gini index measures the probability of a variable being wrongly classified. Hence, a Gini index = 0 is ideal. We selected for finer parametric ranges with Gini index = 0 to further maximize the probability of achieving high evenness. The same process was repeated on the same cluster to discover clusters of high stability within. This process significantly reduced the vast design space into smaller regions that can be comprehensively explored and analyzed.

### Cooperation offers highest flexibility in ecological engineering

We aim to explore how the parametric space varies in terms of evenness as measured by Pielou’s evenness index ^49, 50^ calculated when the system reaches steady state. Pielou’s index measures diversity and species richness. It can vary between 0 and 1, with the former implying that only one species is growing at steady state (no evenness) and the latter meaning that both species have the same biomass in steady state (complete evenness).

Figure 3A illustrates the distribution of evenness in the three models. Cooperation shows the highest evenness with most of the points lying in the space corresponding to an evenness index of 1. The trend is reversed for competition and predation where we noted majority of the points coding for evenness factor 0, indicating higher probability of one species overtaking the other at steady state. These results are significant as they denote that using a cooperation-based co-culture has highest chance of obtaining a highly even system. We observed that predation system had a very binary distribution of evenness. This means that either the system tends to have highest evenness (evenness index = 1) or lowest (evenness index = 0), with very few points lying in the middle, and majority of the points concentrated in evenness = 0 region. Due to the oscillatory behaviour of predator-prey interaction, we expected low evenness due to differences in the biomass of two species. Interestingly, there are regions where the system displays high evenness. This was attributed to specific parameter value combinations that weaken the behaviour of the predator, balancing the interaction to mimic cooperative behaviour (see figure in Supplementary). Therefore, there is scope for predation to have high evenness by changing into cooperation.

**Figure 3:**
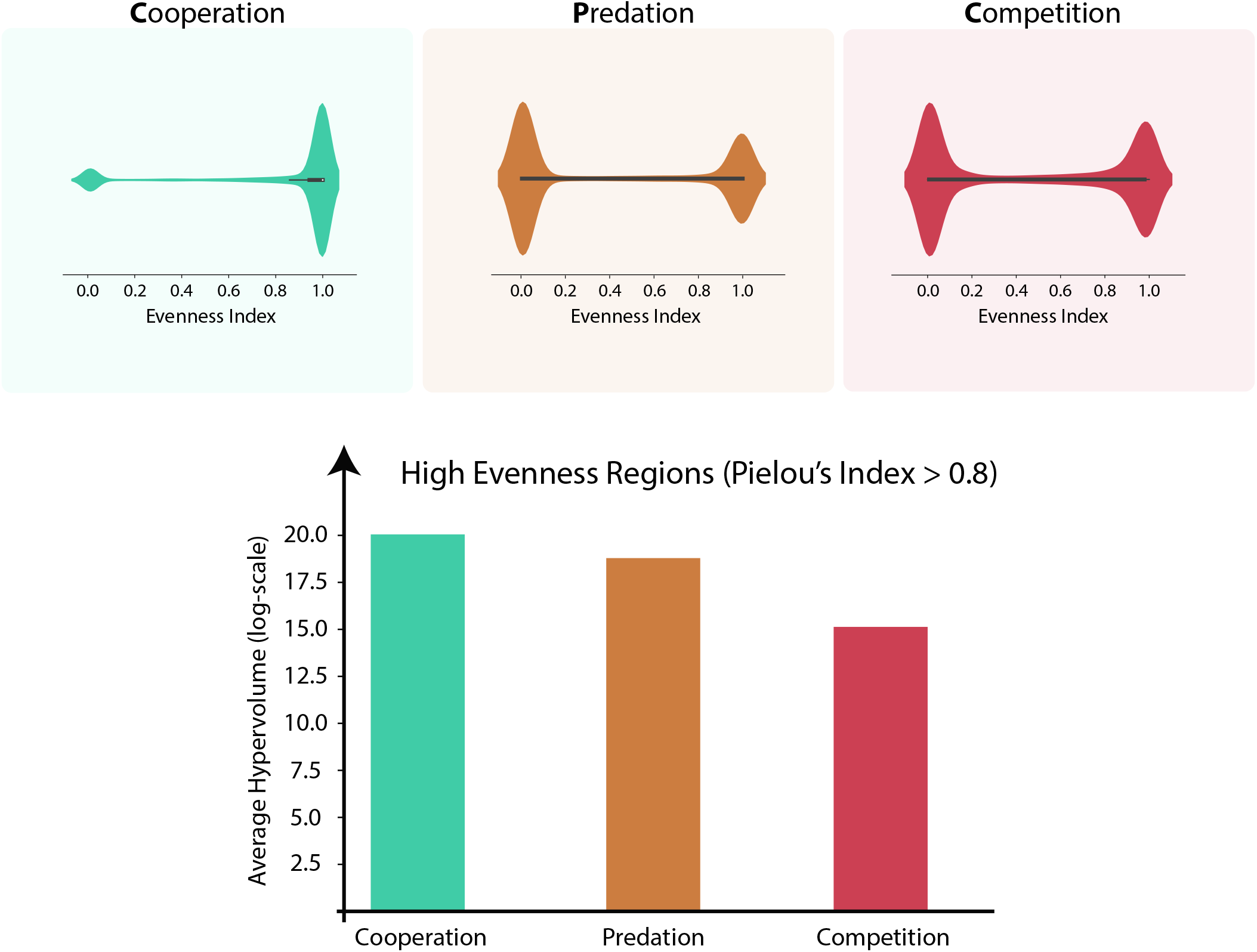
The violin plots show the distribution of evenness in the three models. Cooperation has the highest density of points with evenness index = 1, leading the system to mostly exhibit high evenness. In predation and competition, the highest density is observed for low evenness. Below the violin plots, all high evenness clusters were plotted with the hypervolumes shown on the axis below. Cooperation has clusters of the largest hypervolumes suggesting that greater region of operation. High evenness clusters in competition are the smallest size, implying least amount of flexibility.

The concept of “hypervolume” is typically used to describe ecological niche ^51^. We have introduced it here as the multiplication product of ranges of all the parameters. Given that there are a high number of parameters, the volume calculated is multidimensional. The hypervolume is a direct measurement of the operating ranges to get desired evenness index, with a higher hypervolume implying a larger operating range. Every cluster generated from our method has an evenness index and a hypervolume. We selected clusters of high evenness (evenness index *>* 0.8) in each model, calculated the average volume and have shown them in Figure 3A. For cooperation, the mean hypervolume of high evenness clusters are larger than predation and competition. Meanwhile, competition has the smallest hypervolume by several magnitudes. This indicates that the operating ranges for high evenness in competition are extremely restricted. Cooperation demonstrates wider parametric ranges for high evenness, offering greater flexibility in design of experiment.

### Deconstructing evenness in synthetic microbial consortia

Our framework generates data to provide a roadmap for guiding experimental decisions when building a synthetic microbial consortia with desired properties. This can be achieved by conducting a comprehensive assessment of the vast model spaces. We split our data into high evenness (Pielou’s index *>* 0.8) and low evenness (Pielou’s index *<* 0.3) regions and then extracted the values of the parameters to obtain any trends. Figure 4 demonstrates the distribution of normalized values of parameters for high evenness and low evenness in all three models.

**Figure 4:**
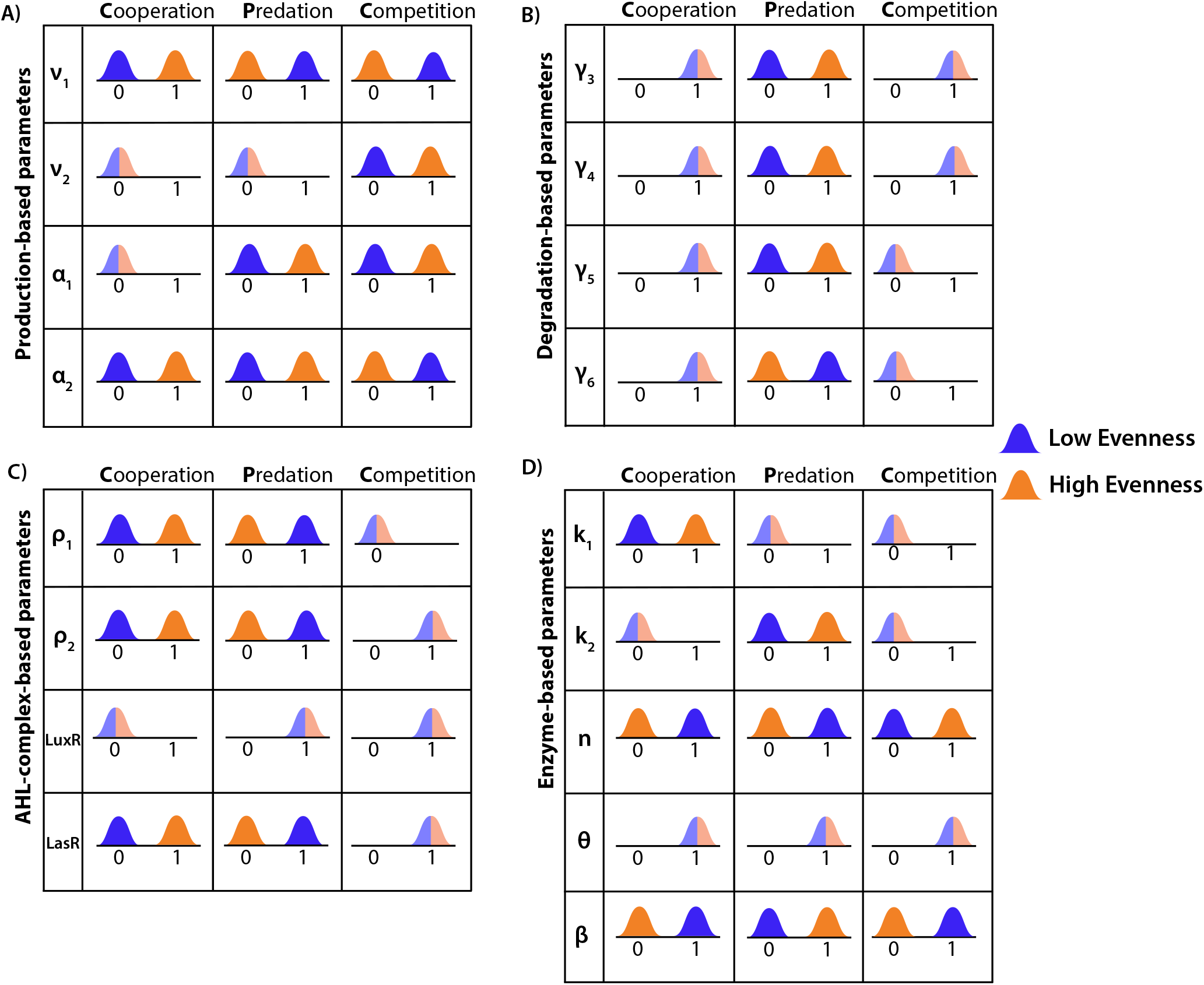
The distribution of the majority of the normalized parameter values for three circuit models in high evenness and low evenness conditions. The parameters were categorized into 4 groups: Production-based (A), Degradation-based (B), AHL-complex-based (C), and Enzyme activity-based (D). It was observed that values of *ν*_1_, *α*_2_, n, *β* parameters are the only ones that change in all three models. In cooperation, *ν*_2_ and *α*_2_ increase to higher values in high evenness, increasing histidine production in histidine-user. In predation, the increase in evenness is marked by the weakening of predator’s actions. For example, *ν*_1_ decreases, *γ*_3_ and *γ*_5_ increase. Meanwhile, the production of histidine in the prey increases with increase in *α*_2_, k_2_, and *β*. In competition model, tryptophan and histidine production is downregulated by decrease in *α*_2_, *β*, and increase in *ν*_2_.

In cooperation model (Figure 4A) exhibiting low evenness, majority of the production-related parameters have lower values, such as *ν, α*, and LasR. This pattern amplifies the difference between the biomasses of the two species particularly by impacting the growth of histidine user. When the model shifts to a high evenness mode, AHL production by tryptophan utilizing species, *ν*_1_ and histidine production rate, *α*_2_ increase. These changes in particular boost the production of histidine and eventually of histidine user, bridging the difference between the two species.

As described before, high evenness cases in predation is attributed to the system mimicking cooperative behaviour. We hypothesized that this becomes possible due to the predator weakening. In our model, the predator is the tryptophan user and prey is histidine user. In Figure 4B, we see predator-associated parameters change to weaken its action on the prey. In high evenness, *ν*_1_ has lower values. The degradation rates also increase, except for *γ*_6_, which is the degradation rate of histidine production in the prey. The prey is further strengthened as *α*_2_ increases. Decrease in *ρ*_1_ coupled with increase in *γ*_3_ decreases the concentrations of AHL1-LuxR complex also weakening the predator’s inhibitory effect on the prey. Moreover, in high evenness the values of the dissociation constant associated with the prey (*β*) increases. These trends illustrate how by weakening the predator and strengthening the prey, a cooperative behaviour is achieved and hence high evenness.

Our results suggest that in competition, transition into high evenness is associated with reducing tryptophan and histidine production in respective species. In high evenness, by lowering histidine production rate, *α*_2_, and dissociation constant of histidine, *β*, the histidine-user decreases histidine production, while an increment in AHL production by histidine-user species, *ν*_2_, strengthens the inhibition of tryptophan production. In competition, coexistence is achieved when each species inhibits its own population before that of the competitor. This is visible when lower but similar levels of biomasses are observed in competition case.Cooperation and competition are symmetrical interactions. However due to the species being auxotrophic for different amino acids, there is a metabolic imbalance as the contribution of tryptophan and histidine to biomass production is different. Since histidine is required in higher amounts for cellular growth, the histidine-utilizing species has a disadvantage in the consortia, having more work to do. In cooperation, it appears that the circuit parameters need to be modulated to compensate for this discrepancy and inject more resources to the histidine user. The patterns that emerge from our results hint that evenness can be manipulated by the concentrating on the species in the consortia that has the greatest resource requirement making it a “limiting” species. It is important to note that the distribution shows where most of the points are and not the entire distribution.

Next we wanted to explore how much the value of each parameter varied between the models. Figure 10 in the supplementary section highlights the normalized variances of each parameter in high evenness regions. Parametric variances deconstruct the hypervolume, determining which parameters are changing to what extent, giving architectural information about the design space of high evenness. We found that each parameter within cooperation showed most variance. This observation was in line with cooperation having the largest hypervolume, as remarkable changes in the parameter values mean broader ranges. The results differed for predation and competition, which displayed lesser general variance, hinting that the design space of those consisted of more limited parameter values.

### Diversity-Stability link is context-driven

Diversity-Stability debate has been central in the field of ecology ^52^, with the relationship between evenness-stability even more unclear. As we move to designing synthetic consortia, this relationship continues to be of utmost importance. Diversity can be decomposed into species richness and evenness. Even though our work focuses on microbial evenness, the use of Pielou’s evenness index which takes into account species richness ensures that the metric is a reasonable reflection of the diversity as well. Furthermore, in our case study we are fixing the species “richness” (number of species in community) to two and only varying relative abundances, therefore the biodiversity of the community is solely dictated by evenness.

In order to characterize the stability of our models, we used linear stability analysis. In summary, the variables are perturbed and Jacobian matrix is computed. The eigenvalues in the matrix are the eigenvalues of the equilibrium and have been used to measure the stability of equilibrium ^53, 54, 55^. We define stability as the tendency of the system to go back to its initial state after perturbation. Figure 5 illustrates the stability of the three models and their relationship with evenness.

**Figure 5:**
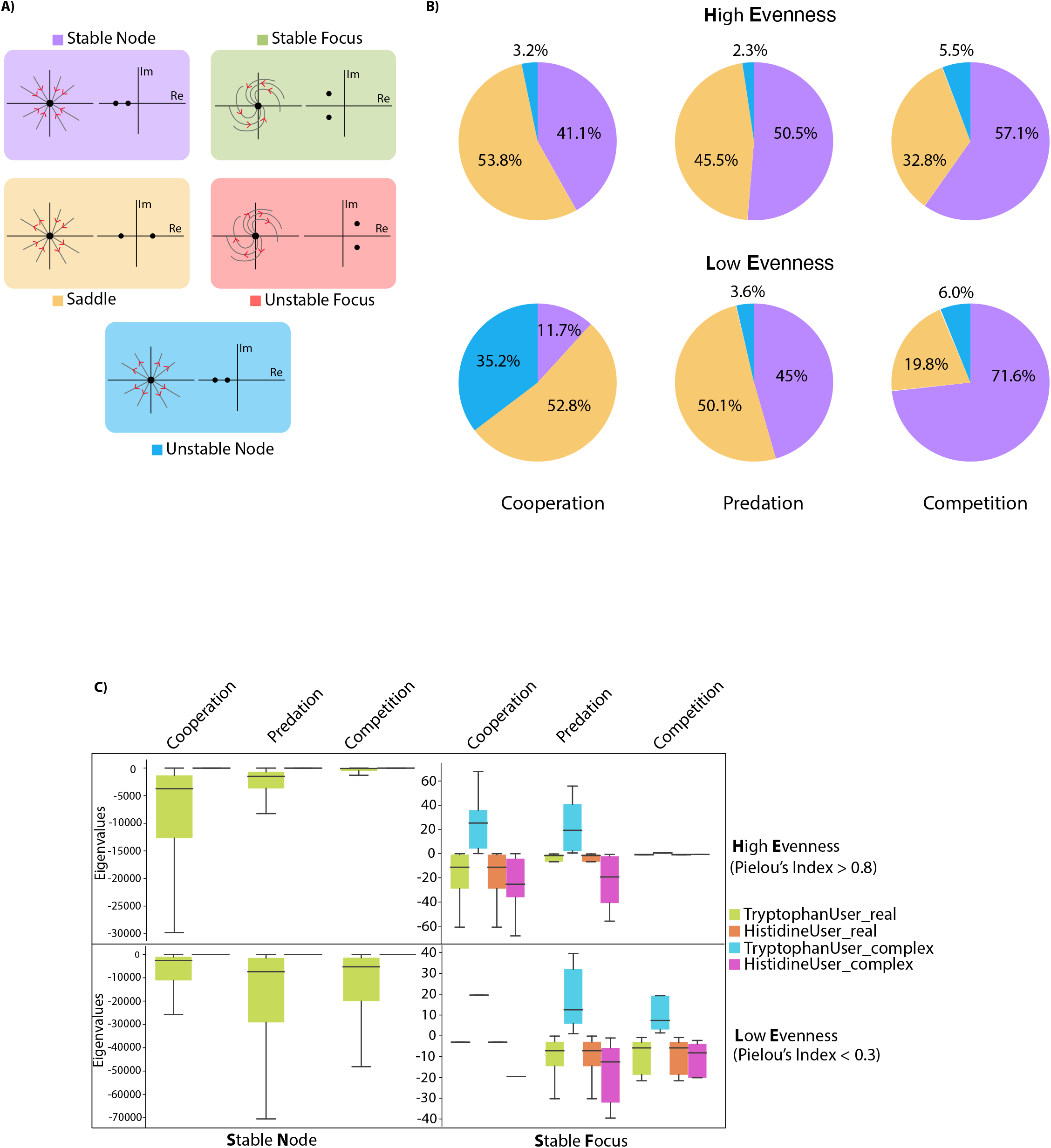
Stability Analysis: Stability was divided into stable node, stable focus, saddle, unstable focus, and unstable node. Using linear stability analysis, eigenvalues were obtained to test the stability of biomass of tryptophan user and histidine user. B) The proportion of each type of stability was evaluated for all models to check how they vary in high evenness (Pielou’s Index *>* 0.8) and low evenness (Pielou’s Index *<* 0.3) C) The eigenvalue distribution of tryptophan user and histidine user are shown for stable nodes and stable focus to assess the link between the frequency of return to steady state versus evenness.

The three synthetic consortia have non-linear dynamics over vast parametric space. Therefore, it is not feasible to assume that the system would have a “single” stable state. Different regions can have different stabilities depending on the parameter values. We treat our system as two-dimensional, as we are interested in the stability of the biomass of both species. For a deeper analysis, we have decoupled stability into five categories: stable node, stable focus, saddle, unstable focus and unstable node. A system is deemed as stable if it has stable node and stable focus. Unstable system is defined by saddle and unstable node and focus points as depicted in Figure 5A. In order to decipher the relationship between evenness and stability, we divided the points into high evenness (Pielou’s index *>* 0.8) and low evenness (Pielou’s index *<* 0.3) and sought to analyze the stability within the two categories. We divided this analysis into parts. First, we inspected the size of stable and unstable regions for our models in different diversity conditions. Second, we investigated the eigenvalue distribution of the two species to determine response rate of the system to a disturbance. Across different consortia in both conditions, majority of the regions fall under stable nodes and saddle points, with stable focus and unstable focus occurring at negligible points. However the proportions vary in relation with evenness. In Figure 5B, we observed that in cooperation, evenness and stability have a positive correlation. In high evenness, stable nodes and saddle points are almost evenly split with the unstable region being slightly larger. However in low evenness, though the saddle points do not change significantly, there is a remarkable increase in unstable node accompanied by a sharp decrease in stable nodes. In contrast, competition displayed the opposite behaviour. Stability and evenness have a negative relationship. When evenness decreases, the region occupied by stable node significantly increases and the saddle points decrease. Predation did not present with a strong link. Much like cooperation, it appears that stability decreases in low evenness, however the change is not by a significant margin. This implies in predation the link between stability and evenness might be harder to substantiate or be non-existent.

We noted that stable regions were not distinctly isolated from unstable regions. For all models, both stable and unstable regions co-exist together, even though the ratios vary with evenness. This suggests that though a pattern can be concluded between evenness and stability for different social ecological interactions, the relationship does not universally hold true. For example, the likelihood of encountering a stable region in high evenness competition is higher, but instability could still be encountered, further complicating the debate between evenness and stability and justifying the need for more intelligent engineering strategies.

The eigenvalue distribution for stable nodes and stable focus are captured in Figure 5C. Larger eigenvalue indicate a faster return to steady state following a perturbation and vice-versa. In high evenness, cooperation and predation co-cultures return to steady state faster than competition. Within high evenness, competition had larger regions of stability compared to cooperation, but its eigenvalues are lower. This signifies that in high evenness conditions species in competition return to steady state slower than in cooperation despite the chances of encountering stable regions being greater in competition. On the other hand, in low evenness, the eigenvalues for competition witness a dramatic increase with predation eigenvalues decreases. In cooperation, the distribution of eigenvalues are not related to evenness. The values for stable focus were also demonstrated even though they do not occur frequently. In cooperation, as evenness decreases so does the eigenvalue distribution of stable focus. Whereas the opposite trend is observed for competition.

The eigenvalues of stable nodes of tryptophan-user biomass is always consistently more negative than that of histidine user despite different ecological dynamics and evenness. It appears that stable node is driven significantly by only one species in the consortia. This difference in eigenvalue was not noted for stable focus. Figure 12 (in supplementary section) shows the eigenvalues of the two species for saddle points and unstable focus corroborating the pattern noticed for stable node. This was a curious finding we attributed to the difference in the growth rates of the species, which is being controlled by the system to bring high evenness conditions. It appears that the faster growing species in a community is dampened to achieve stability. We believe this is an important insight into stability. When designing synthetic consortia, the species should be chosen and/or engineered to account for this.

Our analysis proves that the evenness-stability relationship is complex and contextual. Additional challenges are imposed with cases like predation where the association between the two is not as clearly defined. Furthermore in synthetic microbial community comprised of multiple species and hence multiple social interactions, the relationships are expected to change. Therefore, there is a need to regulate stability to prevent dependence on a variable qualitative relationship.

### Predicting optimal ranges for high evenness and high stability

An ideal synthetic microbial consortia would be highly even and stable to ensure long term function and survival. Using our workflow, we aimed to engineer the synthetic consortia to operate in both high evenness and high stability regions. As per our process, we first partitioned the design space into clusters of high evenness, following which we selected a suitable cluster with highest probability and appropriate hypervolume. More samples were generated within the chosen cluster using sobol sequence for analysis. Next, we evaluated the stability at each point and obtained ranges by selecting for stable points and eliminating unstable focus, nodes and saddle points. This method yielded multiple clusters from which we chose a stable region. The even and stable operating ranges have been shown in Figure 6. The parameter values should be set within the darker shaded region to maximize the probability of the system exhibiting high evenness and high stability. The operating ranges for cooperation highlighted are very wide. This is a positive trait as it gives scientists more flexibility when engineering the circuit parameters. The optimal ranges became narrower for predation and competition, making circuit tuning more challenging. Figure 13 in supplementary information illustrates the average hypervolume of the high stability regions within high evenness. Predation is shown to have the smallest hypervolume, suggesting that the operating ranges to achieve high stability are the narrowest. However as seen in Figure 3, competition has the smallest hypervolume for high evenness, not predation. This shows that the hypervolumes for high evenness and high stability are not correlated.

**Figure 6:**
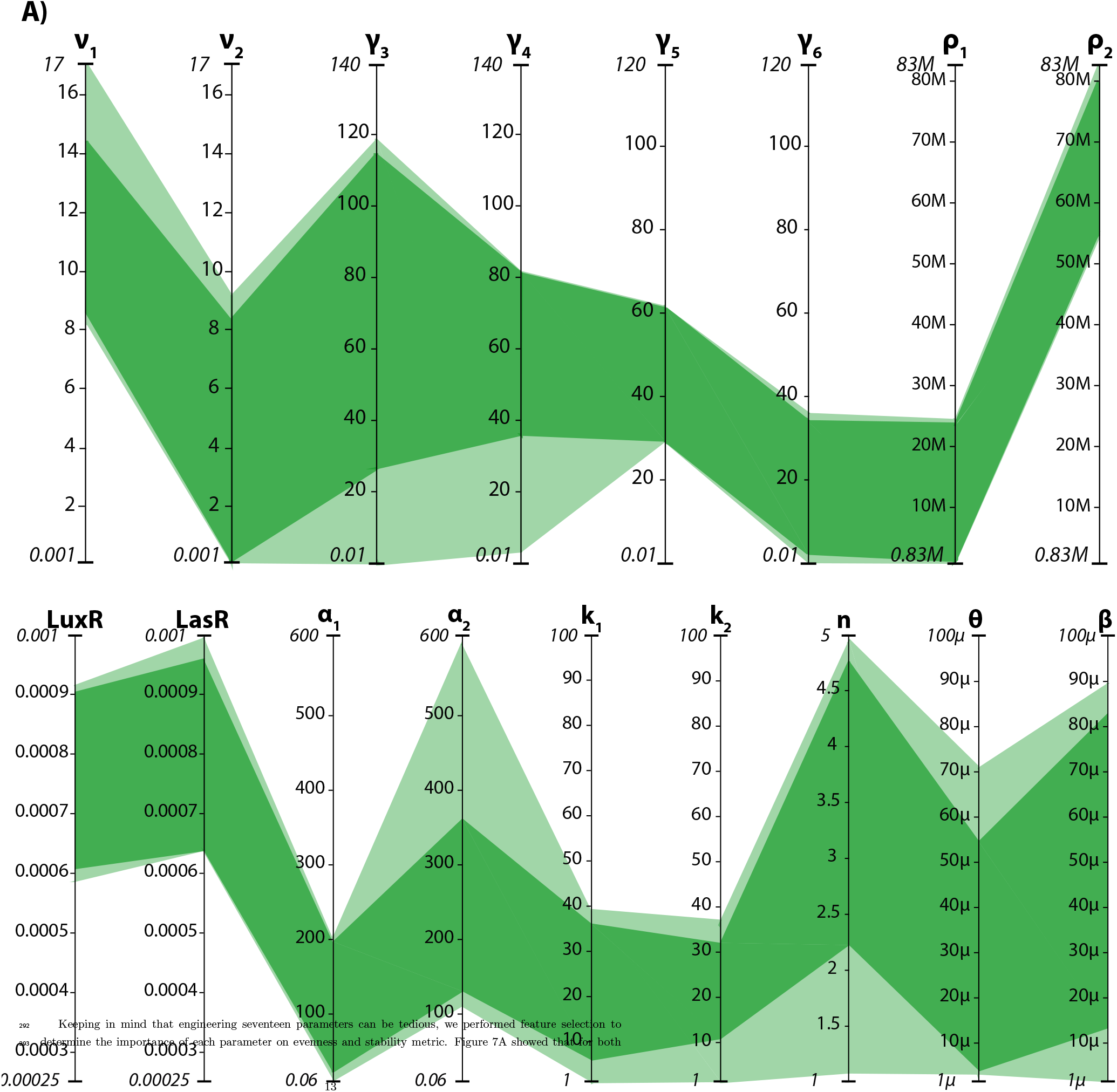
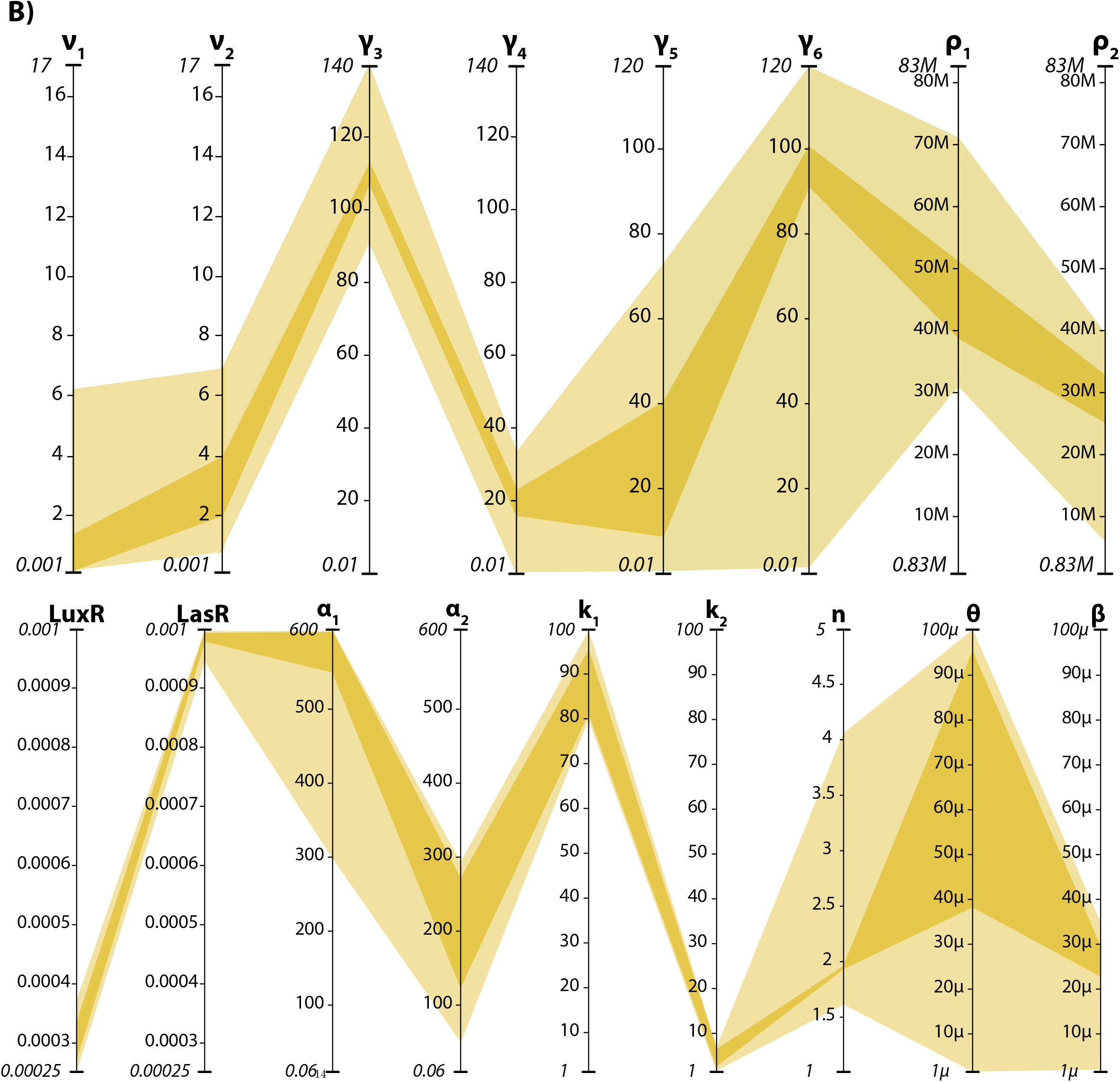
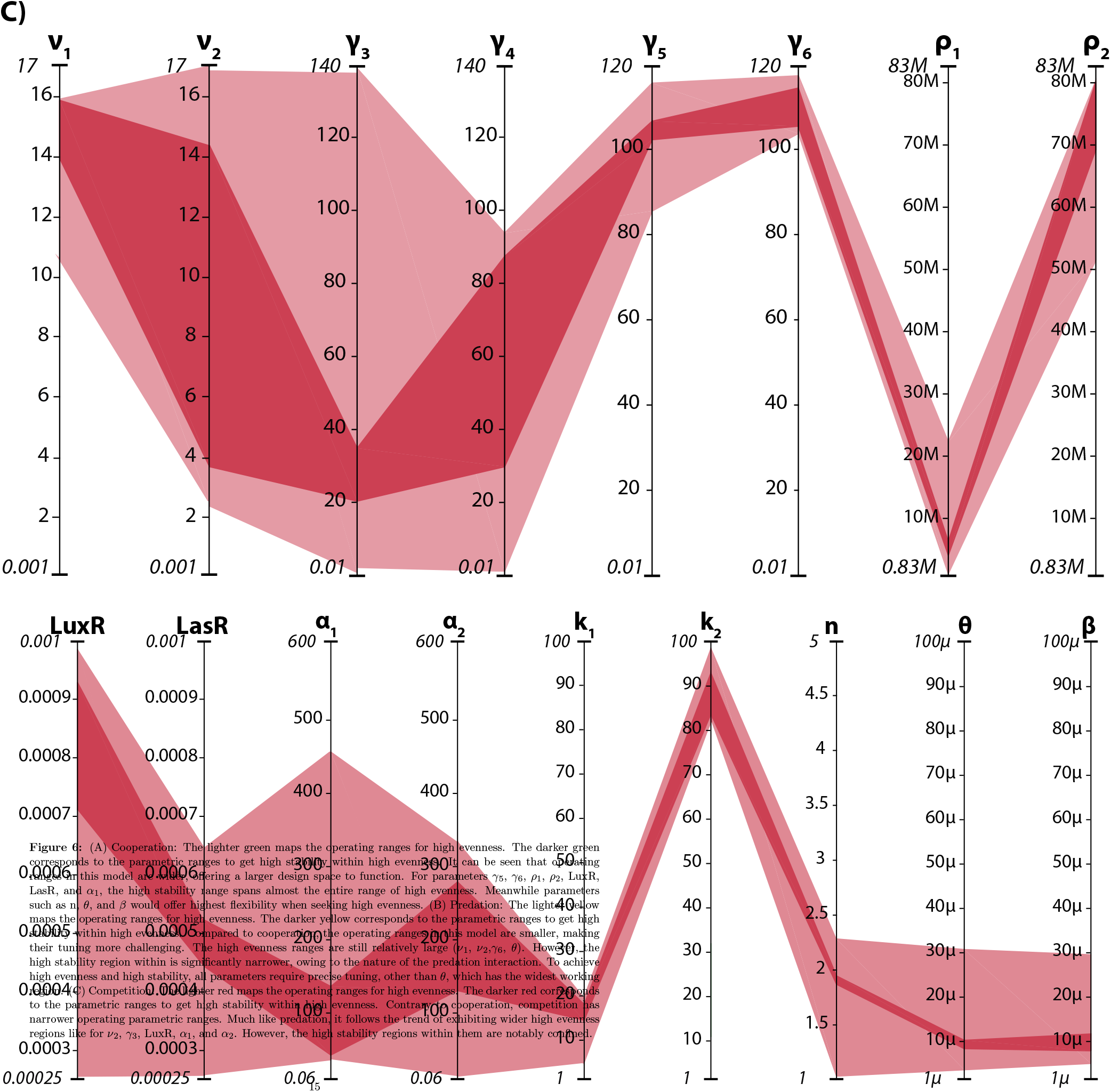
(A) Cooperation: The lighter green maps the operating ranges for high evenness. The darker green corresponds to the parametric ranges to get high stability within high evenness. It can be seen that operating ranges in this model are wider, offering a larger design space to function. For parameters *γ*_5_, *γ*_6_, *ρ*_1_, *ρ*_2_, LuxR, LasR, and *α*_1_, the high stability range spans almost the entire range of high evenness. Meanwhile parameters such as n, *θ*, and *β* would offer highest flexibility when seeking high evenness. (B) Predation: The lighter yellow maps the operating ranges for high evenness. The darker yellow corresponds to the parametric ranges to get high stability within high evenness. Compared to cooperation, the operating ranges in this model are smaller, making their tuning more challenging. The high evenness ranges are still relatively large (*ν*_1_, *ν*_2_,*γ*_6_, *θ*). However, the high stability region within is significantly narrower, owing to the nature of the predation interaction. To achieve high evenness and high stability, all parameters require precise tuning, other than *θ*, which has the widest working region. (C) Competition: The lighter red maps the operating ranges for high evenness. The darker red corresponds to the parametric ranges to get high stability within high evenness. Contrary to cooperation, competition has narrower operating parametric ranges. Much like predation, it follows the trend of exhibiting wider high evenness regions like for *ν*_2_, *γ*_3_, LuxR, *α*_1_, and *α*_2_. However, the high stability regions within them are notably confined.

Keeping in mind that engineering seventeen parameters can be tedious, we performed feature selection to determine the importance of each parameter on evenness and stability metric. Figure 7A showed that for both evenness and stability, *ν*_2_ was the highest contributing factor in cooperation. Though in evenness, cooperativity and dissociation constant appears to be important as well, it is in stability we notice that several parameters are significant contributors, in particular degradation rates and protein production rates. Manipulating them in a wet-lab setting is straightforward by changing the RBS strengths ^56^. In predation, the operating ranges become noticeably narrower, particularly for high stability within high evenness. However feature selection in Figure 7B shows that *ν*_1_ dictates evenness and stability in predation model. The rest of the parameters seem to have a markedly lower to negligible impact. We believe that lowering *ν*_1_ has a cascade effect of weakening the effect of the predator on the prey, thereby pushing the system toward high evenness and stability. In competition, we note narrow engineering space in Figure 6 and feature importance shows multiple parameters being of significance. Most important parameters shared between evenness and stability are n, LuxR, *α*_2_, out of which the first two have a very limited tuning range according to the cluster we have chosen but can be engineered experimentally^57^. This insinuates that competition model is harder to engineer, requiring very specific values.

**Figure 7:**
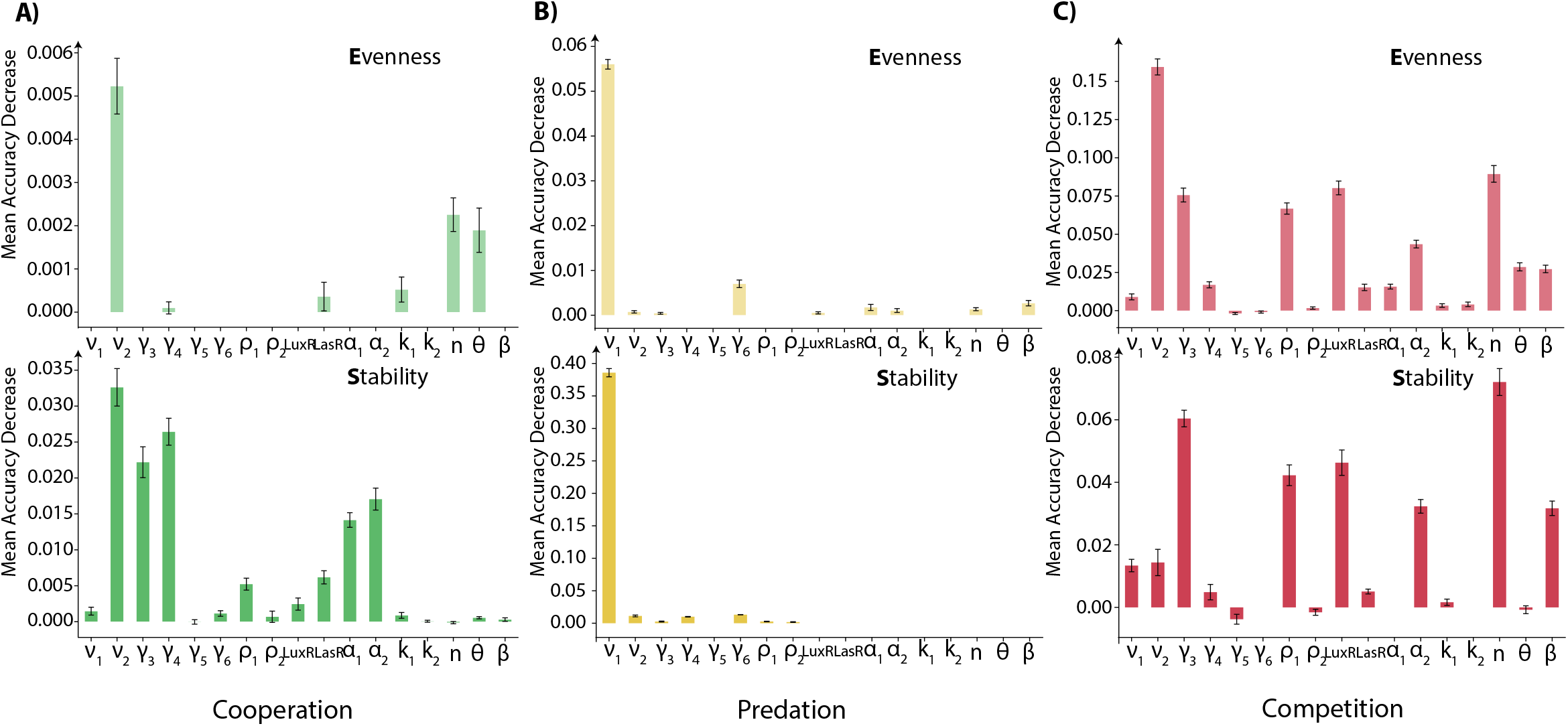
Feature Importance: Circuit parameters were ranked on their contribution toward evenness and stability for all interaction circuits using the mean accuracy decrease method shown on the y-axis. Differences were noted for each. A) Cooperation: *ν*_2_ was the most important parameter for both evenness and stability. For stability, there were more parameters of significance especially degradation rates and protein production rates. There are more parameters important in determining stability than evenness. B) Predation: Both evenness and stability are prominently driven by *ν*_1_ alone. C) Competition: This model witnesses highest number of significant parameters compared to the other two models. For evenness *ν*_2_ is most important and for stability it is n. Competition generally witnesses several parameters contributing to evenness and stability compared to the other two models.

Tuning the circuit parameters to have values that fall within operating ranges maximize the probability of obtaining high evenness and high stability in synthetic microbial consortia. However that output is not fully guaranteed. The engineering decisions can further be aided by the results in Figure 4. For example, for parameters that have a very wide operating ranges, looking at the general distribution for high evenness in Figure 4 can guide which value to select. This is particularly useful for cooperation where the flexibility is very large. Large engineering space is desirable, however it would also contain the most fail-points. Therefore, there is a tradeoff between accuracy and size of the design space. Our computational framework incorporates another step that can help eliminate fail-points. Applying decision tree on the selected clusters will eliminate a portion of those fail-points. Figure 11 in supplementary information shows decision tree for cooperation and predation high evenness clusters and the further circuit tuning that needs to be conducted to increase the probability of getting high evenness.

## Discussion

Diversity and stability are essential properties of microbial communities, however the relationship between them remains a subject of interest. Historically, the studies have concentrated on the macroscopic realm where the general consensus has been a positive relationship. However, evidence has pointed to a negative relationship and in some cases no relationship at all. Recently, research has been expanded to micro-ecosystems. While most data demonstrate a positive diversity-stability link, the relationship is not universally applicable as the problem is intricate with different conditions giving rise to different interplay^58^. Studies have characterized this link in terms of strength of interaction, types and frequency of interactions, nutrient availability, and food-web topology ^59, 60, 61, 55^. Diversity is defined by richness and evenness. In this work, we have focused on evenness, which remains an extremely understudied topic in this field. We underline that context matters when trying to answer this question.

In our paper, we have demonstrated evenness-stability relationships based on mode of interaction for synthetic co-cultures. Furthermore, we have decomposed stability for each model to gain a deeper understanding into what comprises their stability. Though research into effectively characterize this relationship is mounting, to our knowledge there is no study that aims to engineer this link. As opposed to viewing evenness-stability as intrinsic properties of a synthetic consortia, we propose that they are mutable targets. By engineering parameters, we can shift a synthetic consortia to a region of high evenness and high stability. Arming synthetic biologists and ecologists with this ability, increases their freedom of choosing synthetic models, as evenness and stability are no longer fixed restrictions. This is in particular useful, as the mechanisms behind stability remain elusive.

Diversity is considered to be one of the factors driving stability, but other factors such as substrate storage, habitat characteristics, biotic and abiotic interactions, and environmental conditions are also contributors. Variations between different synthetic consortia can hinder the establishment of a universal correlation with different mechanism yielding different relationships. Though a comprehensive understanding of stability will be an incredibly useful insight, our methodology proves that high stability regions can be identified without that knowledge. We can directly engineer the system to have high stability without changing other factors. Those factors can be accounted for but our method can easily be modified to characterize any parametric space and find a solution. We have demonstrated the concept for co-cultures, but it can easily be expanded to multiple species with a combination of interactions with parameter values that narrow the function in a high evenness and high stability region.

We emphasize that while some relations emerge, the parametric space is too vast and multi-dimensional to find design rules that hold universally true. In fact, our analysis hints that even generally held true relationships can shift and modify depending on the conditions. Therefore when designing optimal functioning space for synthetic consortia, it is important to not rely on general relationships as they are several fail points. Our strategy of narrowing down parametric spaces and selecting suitable clusters to operate within will alleviate this issue and maximize the probability of seeing desired effects.

Some research groups have attempted to survey the design space of genetic circuits ^62, 63, 64^. Our approach differs as we use adaptive sampling to cover the design space and surrogate model to check the coverage. The commonly used method to explore design space has been uniform sampling which has a major drawback. There is an extremely high computational cost, making it an unsuitable technique to study complex and vast spaces. A common strategy has been to set a benchmark of number of simulations to perform, but there is no way to know if the design space was covered efficiently. Our adaptive sampling-based approach combined with principles of machine learning partitions the design space into pockets of high evenness within which can we identify regions of high stability. Even though in context of synthetic consortia, this method can find wider application. For example, in metabolic engineering using co-cultures, this method can be used to optimize the system to maximize chemical production. The method can be broadly applied to explore and evaluate any genetic circuit. Additionally, the ability deconstruct solution space based on multiple metric will prove particularly profitable to synthetic biologists.

Designing synthetic consortia is a multi-faceted challenge. One of the approaches involves optimal tuning and balancing the circuit parameters for desired output. Most of the methods to design synthetic microbial consortia center around model selection and configuration design ^33^. We propose an alternative perspective: optimizing synthetic ecological circuits. The main advantage with this approach is the freedom to design models without any structural constraints. Our methodology can easily be adapted to fit any synthetic ecological configuration and enhance its performance by determining regions of desired output. One of the most unique and powerful aspects of our developed method is the comprehensive exploration and analysis of the design space. As opposed to designing the structure of synthetic consortia, we analyze the architecture of the design space of synthetic consortia to extract valuable insights into behaviour and instructions on how to optimally regulate it. We believe our framework will aid not only in ecological engineering but also the diverse field of synthetic biology.

## Methods

### Models and Packages

Genome-scale metabolic model of *E*.*coli* iML1515 was obtained from BiGG Models database. All coding was done in Python language.

### System Equations

The following ordinary differential equations are used to describe the circuit dynamics in a chemostat setting. It is assumed that the chemostat is perfect, which means flow rate of media in and out of the reactor is the same, keeping volume constant.

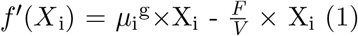

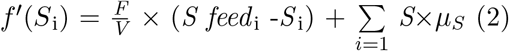

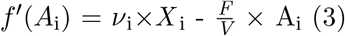

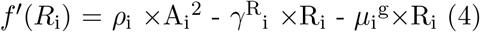

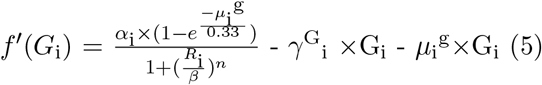

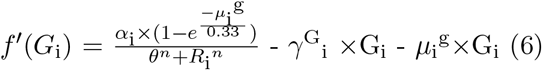

Variables: X = biomass concentration, *μ*^g^ = growth rate calculated by FBA, *F* = flow rate of fresh media, *V* = volume of the media in the reactor, S = substrate concentration, S feed = substrate concentration in the incoming media, *μ*_*S*_ = substrate consumption rate, A = AHL molecule concentration, *ν* = AHL production rate, R = AHL-Protein complex concentration, *ρ* = dimerization constant, *γ*^R^ = degradation rate of complex, G = gene concentration, *α* = protein production rate, *γ*^G^ = protein degradation rate, n = cooperativity, *θ* = Dissociation constant associated with activation, *β* = Dissociation constant associated with repression. 0.33 /hr = growth rate ^65^

### Steady State Error

Biomass of species 1 latest = B_1_L Biomass of species 1 previous = B_1_P

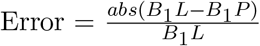

Steady state is detected when the error *<* 1e-3 and when the biomass of both species is above 0.1 to ensure that there was growth. All simulations where washout occurred or where the biomass of the species did not grow were excluded from stability analysis.

### Parameters

| Parameter | Minimum Value | Maximum Value | Unit | Reference |
| --- | --- | --- | --- | --- |
| $\nu$ | 0.001 | 17 | mM hr<sup>-1</sup> | <sup>66</sup> |
| $\gamma^R$ | 0.014 | 140 | hr<sup>-1</sup> | <sup>66 67</sup> |
| $\gamma^G$ | 0.012 | 120 | hr<sup>-1</sup> | Reference |
| $\rho$ | 830000 | 83000000 | mM<sup>-3</sup> hr<sup>-1</sup> | <sup>67</sup> |
| LuxR/LasR | 0.00025 | 0.001 | mM | Bionumbers |
| $\alpha$ | 0.06 | 600 | mM hr<sup>-1</sup> | <sup>67</sup> |
| $k$ | 1 | 100 | Unit | Bionumbers |
| $n$ | 1 | 5 | Unitless | <sup>67</sup> |
| $\theta$ | 0.000001 | 0.0001 | mM<sup>2</sup> | <sup>67</sup> |
| $\beta$ | 0.000001 | 0.0001 | mM<sup>2</sup> | <sup>67</sup> |

### Adaptive Sampling

For implementation, a python package, pymoo - multi-objective optimization in python - was used.

For a target function *f* in *D* ∈ ℝ^n^, initial set of samples are generated **X** = (*x*_*1*_, …, *x*_*m*_). New points, **X**_new_ are added iteratively by solving an bi-optimization score function.

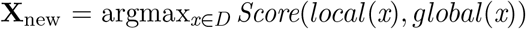

global(*x*) covers the entire domain looking for unexplored regions of interest. Whereas local(*x*) refers to local exploitation, in which the algorithm samples points in regions with high prediction errors. The score is optimized using weights *w*_local_ and *local* + *w*_global_for local exploitation and global exploration respectively:

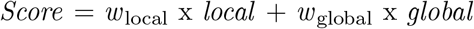

The weights satisfy the criteria: *w*_local_ x *local* + *w*_global_ = 1

In order to find the optimal trade-off between local exploitation and global exploration, ’greedy’ strategy, a form of switch strategy, is implemented. A random value *r* is compared to a threshold value *ε*. If *r* ¡ *ε*, the sampling process switches from local exploitation to global exploration. Using a fixed point, this strategy switches from exploring the entire space to reduce domain uncertainty to local exploitation, which improves model accuracy.

### Surrogate Model Construction

A surrogate model is an ML model trained on adaptive sampling. Our input variables are the parameters whereas as output is the model response we are evaluating: evenness and stability.

An open-source Python package - surrogate modelling toolbox (SMT) was used. During initial training, KPLS^68^ was used, which searches the direction which maximizes variance between input and output variables. As the number of points start increasing, the SVM models^69^ are used as surrogates in clusters, which are also known to be highly effective in high dimensional problems.

### Cluster Selection

We performed density-based clustering, which is recommended for high-dimensional data, to divide our points into high evenness and low evenness clusters. Points with an evenness index above 0.8 was labelled as high evenness. We calculated the hypervolume for each cluster, a product of the parameter ranges that contain points of a specific metric. We identified regions of high evenness within the parametric space and suggest that within these regions, the two species can co-exist as a result of the high evenness index. Additionally, we suggest that the larger hypervolumes of such high evenness regions imply robustness across a range of parameter values.

### Linear Stability Analysis

For an n-dimensional system, x = (x_1_, x_2_,…,x_n_) can be described by ordinary differential equations:

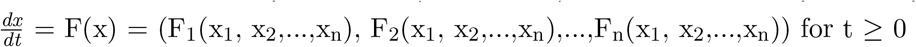

with the initial conditions, x(t = 0) = x_0_ and F(x) is a matrix nonlinear function of x. Following Taylor’s expansion: 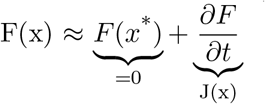

Around fixed points: F(x^*^) = 0

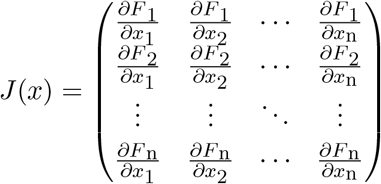

The general solution (close to fixed point x^*^):

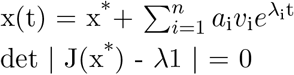

where *λ*_i_ are eigenvalues, v_i_ are eigenvectors and a_i_ are constants determined by using initial conditions

## Code Availability

The code for this study is available on GitHub: https://github.com/LMSE/community_engineering

## Acknowledgements

Authors would like to acknowledge funding from Medicine by Design program at the University of Toronto and NSERC.

## Supplementary Information

**Figure 1:**
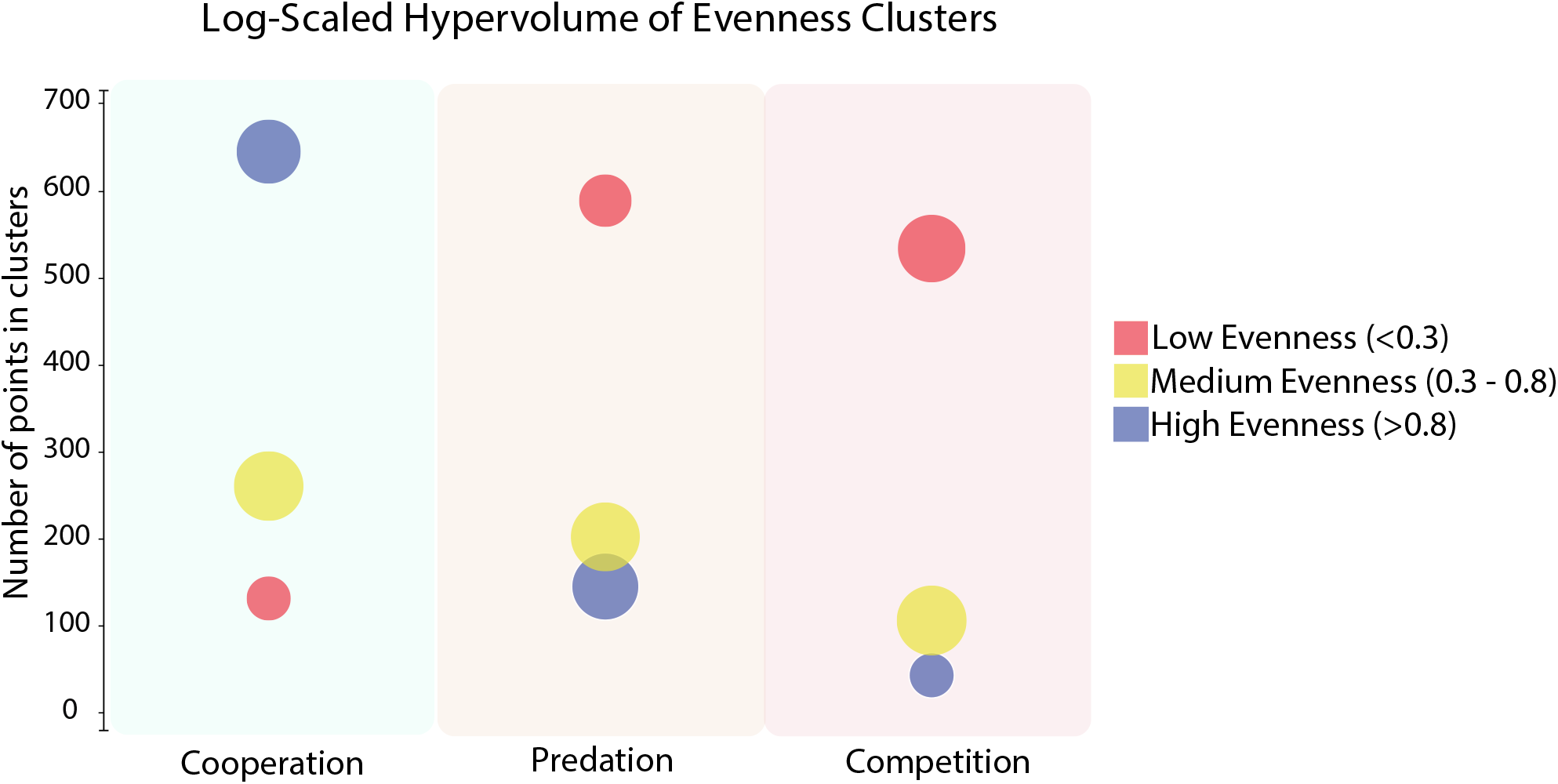
Hypervolume analysis: The entire space was split into low, medium, and high evenness clusters and plotted against the number of points falling inside each cluster. The hypervolume and number of points inside a cluster are not always correlated. As can be seen in predation, the hypervolume of high and medium evenness is larger than low evenness, however the number of points falling within low evenness is the highest. In cooperation, the cluster populated by most points belong to high evenness and in competition the opposite is observed. However, it is in predation that we note that low evenness cluster, which has the highest number of points, has a lower hypervolume compared to high evenness and medium evenness clusters. These results show that the while majority of the points in predation code low evenness, the ranges of the parameters that would yield low evenness are narrowest, reducing the hypervolume.

**Figure 2:**
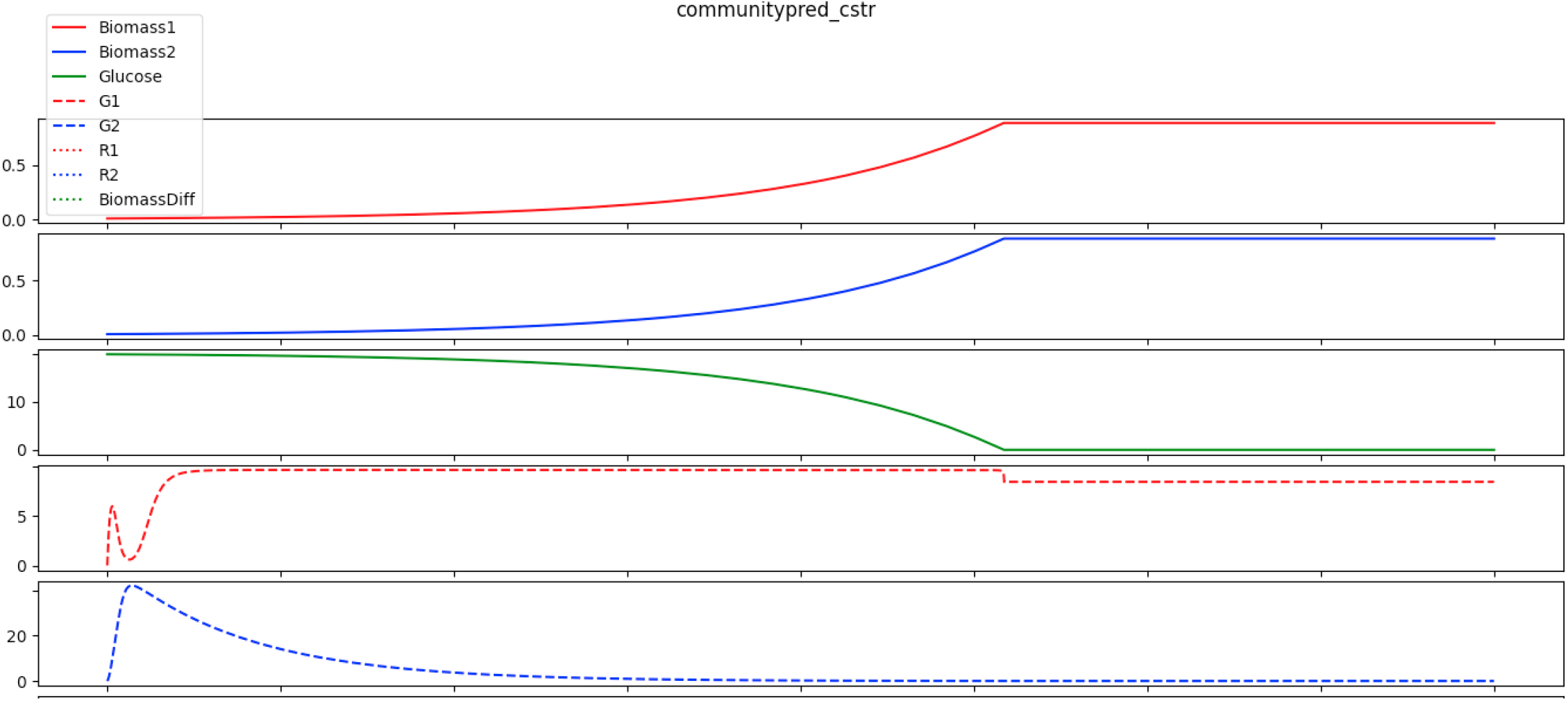
A snapshot of DMMM output for predation after inputting parameter values inside a high evenness cluster. lt can be seen that two species grow together and have the same biomass, showing cooperative behaviour. TRPAS2 (gene of predator/G1) and HISTD (gene of prey/G2) show prey-predator interaction. This proves that it is possible to engineer predation to have high evenness.

**Figure 3:**
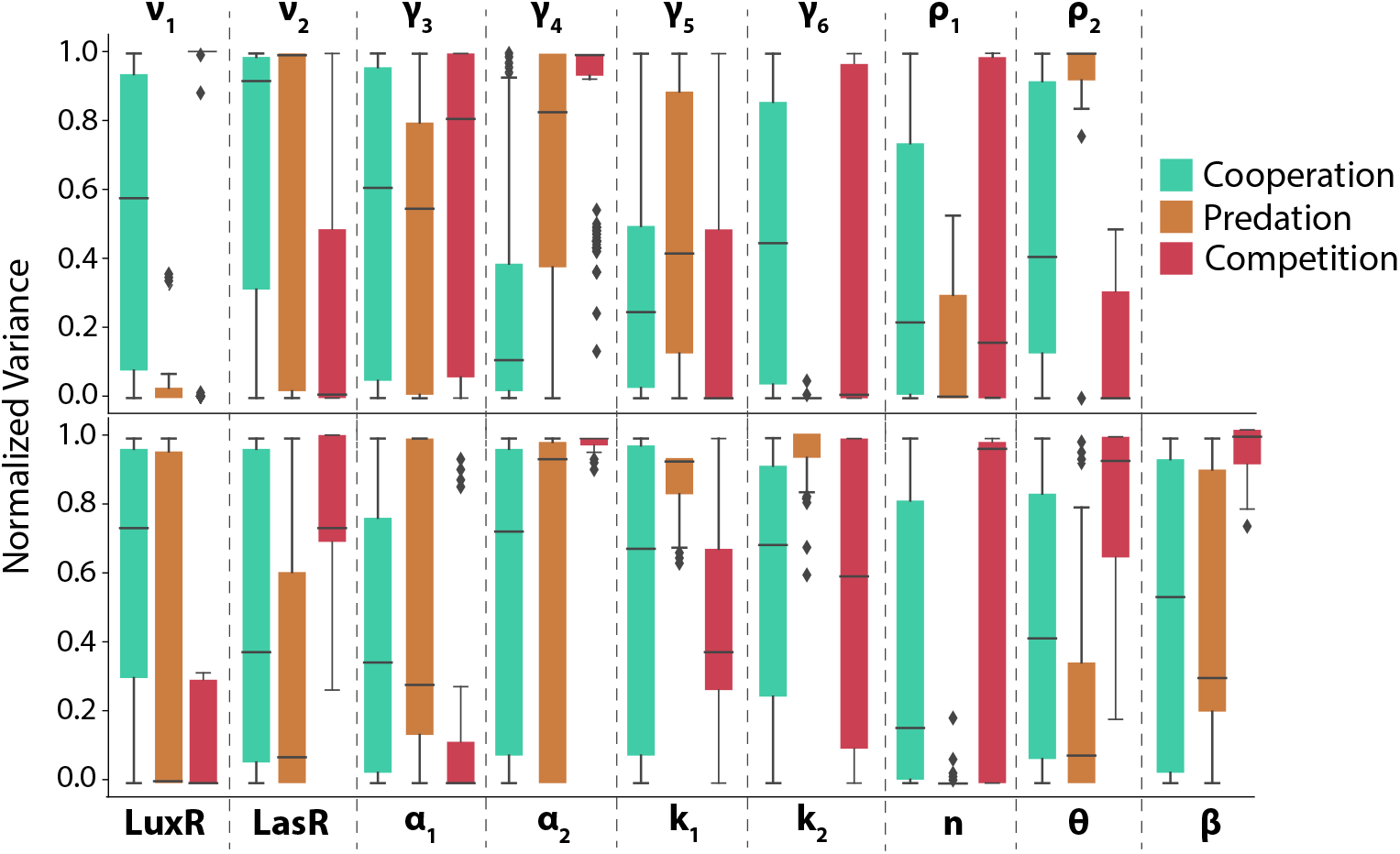
Variance of circuit parameters in high evenness condition gives another angle of how parameters change. Cooperation shows the maximum variance across all parameters.

**Figure 4:**
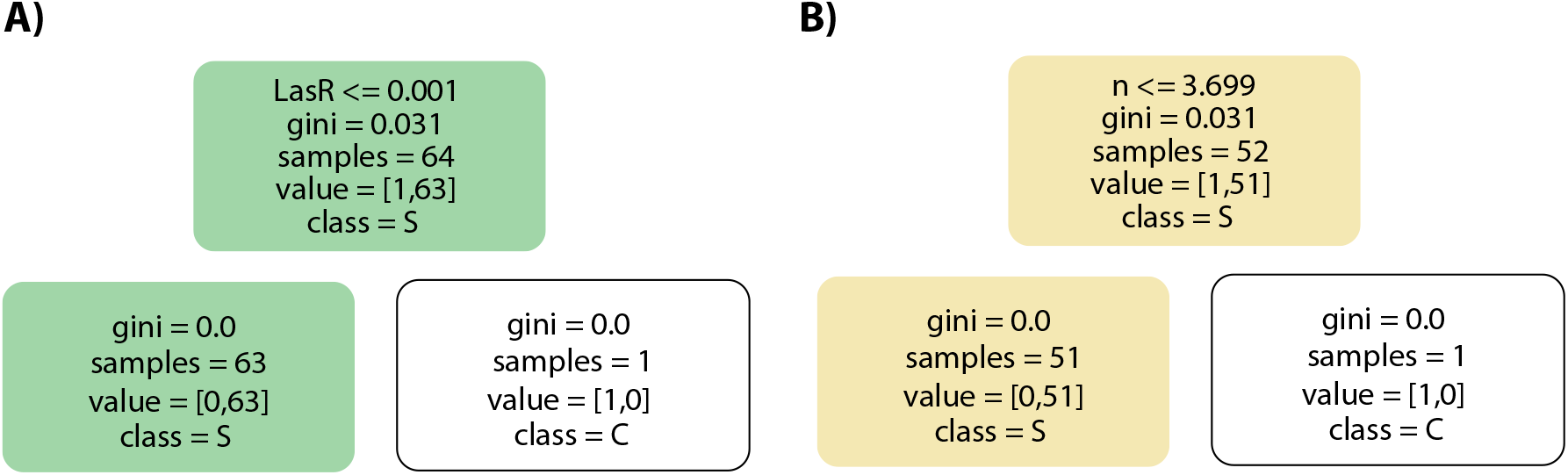
Decision Tree: The evenness clusters to extract the operating ranges from for cooperation and predation had lower probability. Decision tree was applied to identify the parameter that required further tweaking to maximize the probability of getting high evenness.

**Figure 5:**
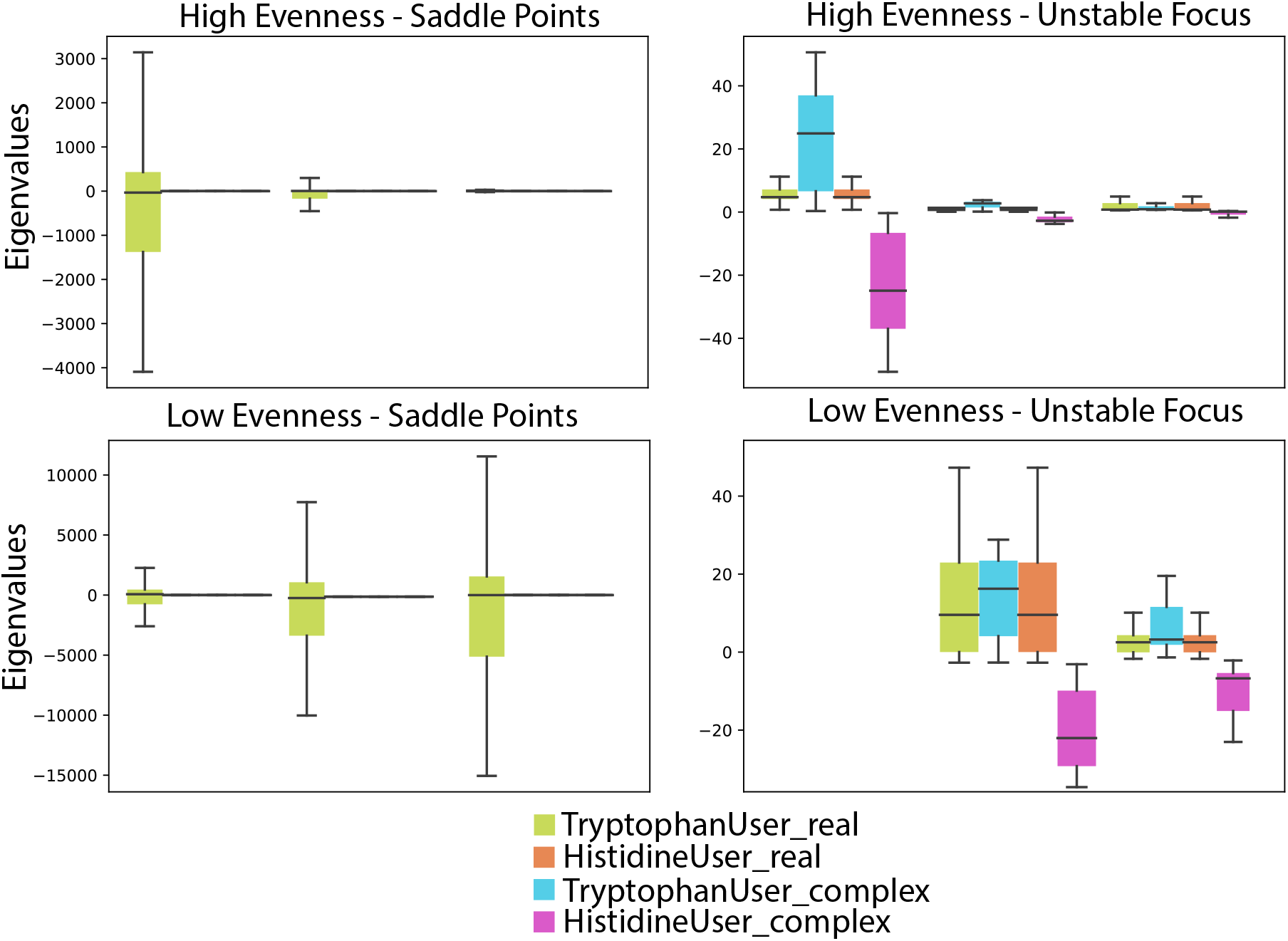
Eigenvalue distribution of stable nodes in high evenness: Results show that the eigenvalues of biomass of tryptophan user are typically larger with greater variation across the three ecological dynamics. This gives an insight into how the metabolic properties of the species affect the stability of a synthetic microbial system

**Figure 6:**
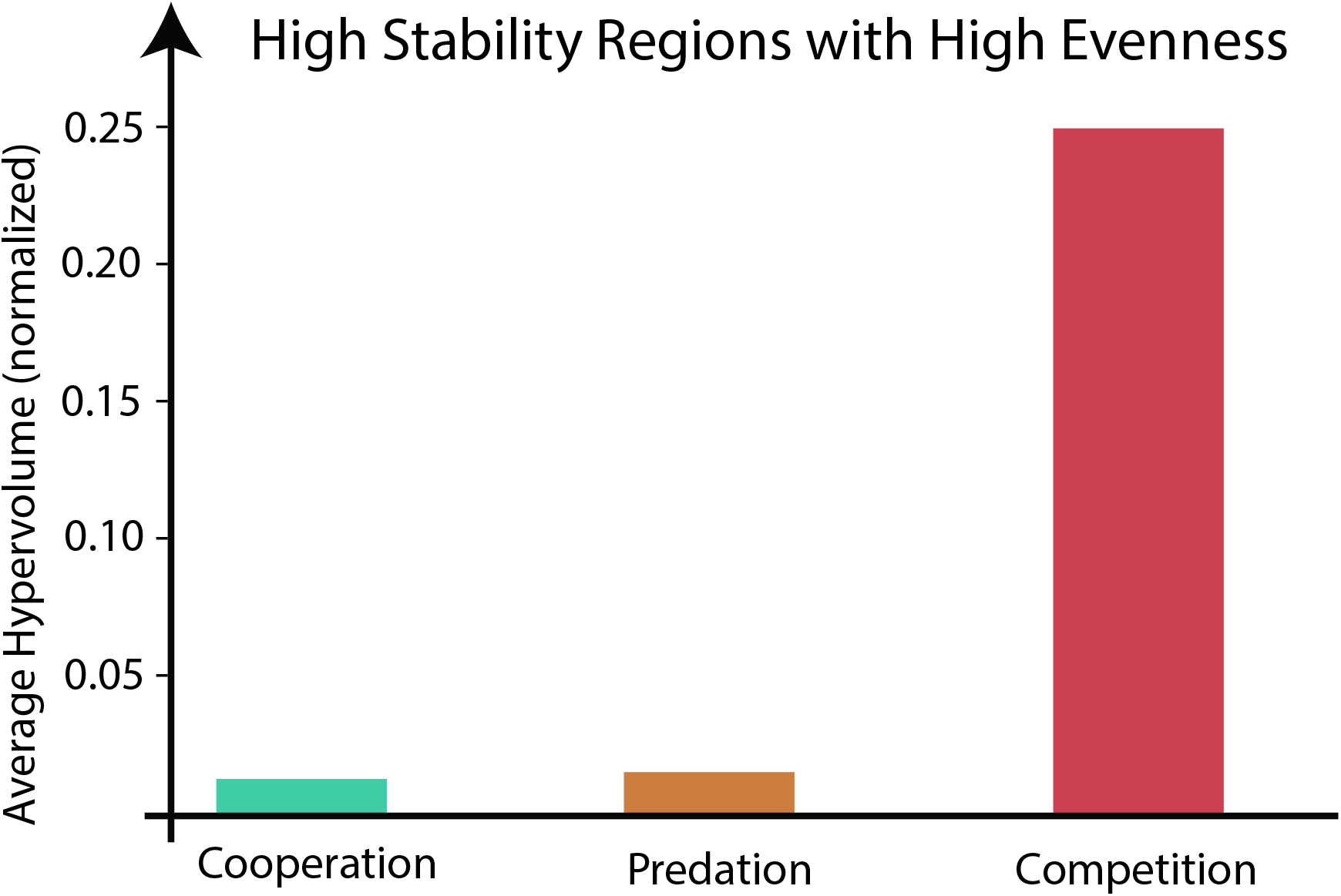
The hypervolumes of the high stability regions within high evenness were averaged. As supported by Figure 6, cooperation has the highest hypervolume, implying that the optimal ranges for high stability are the widest. Predation was shown to have the smallest hypervolume, meaning that the optimal ranges to achieve high stability within high evenness in predation were the most restrictive. Meanwhile, competition has a bigger hypervolume than predation but smaller than cooperation. Even though as seen as in Figure 3, competition has the smallest hypervolume for high evenness but that is not the case for high stability.

